# Molecular tools for phytoplankton monitoring samples

**DOI:** 10.1101/339655

**Authors:** Bárbara Frazão, Alexandra Silva

**Author notes:** Author to whom correspondence should be addressed; E-Mail (F.L.).

## Abstract

HABs can have severe impacts in fisheries or human health by the consumption of contaminated bivalves. Monitoring assessment (quantitative and qualitative identification) of these organisms, is routinely accomplished by microscopic identification and counting of these organisms. Nonetheless, molecular biology techniques are gaining relevance, once these approaches can easily identify phytoplankton organisms at species level and even cell number quantifications. This work tests 12 methods/kits for genomic DNA extraction and seven DNA polymerases to determine which is the best method for routinely use in a common molecular laboratory, for phytoplankton monitoring samples analyses. From our work, Direct PCR master mix for tissue samples, proved to be the most adequate by its velocity of processivity, practicability, reproducibility, sensitiveness and robustness. However, brands such as Omega Biotek, GRISP, Qiagen and MP Biomedicals also showed good results for conventional DNA extraction as well as all the *Taq* brands tested (GRISP, GE Healthcare Life Sciences, ThermoFisher Scientific and Promega). Lugol’s solution, with our tested kits did not show negative interference in DNA amplification. The same can be said about mechanical digestion, with no significant differences among kits with or without this homogenization step.

## 1. Introduction

Phytoplankton are photosynthesizing microscopic organisms that play an important role in primary production, forming the base of the food chain. Under certain ecological conditions, they can flourish and form dense blooms. In the ocean, Harmful Algal Blooms (HABs) also called as “red tides” pose a serious threat to fisheries, public health, tourism, and coastal ecosystems on a pan-global scale (Lee et al. 2013). HABs can cause fish-killing, which can affect economically aquacultures or be accumulated in filter-feeding bivalves (Han et al. 2016) that by itself is not a problem to animals, however can cause illness and death in human consumers.

Phytoplankton samples from long-term monitoring programs or cruises, are preserved with fixatives, such as Lugol’s solution, formalin or glutaraldehyde for subsequent identification of species composition by microscopy (Bertozzini et al. 2005). In classical monitoring programs, samples are fixated with Lugol’s solution, slightly alkaline, at 0,4% (v/v). Conventional light microscopy or phase contrast is the most employed technique in monitoring programs since it is the least expensive, direct and simple method to obtain results of organism’s identification and counting. However, fluorescent techniques and electronic microscopy such as scanning (SEM) or transmission (TEM) can also be used for more detailed analysis, but not as a routine technique.

From some years now, molecular biology procedures are gaining expression in monitoring programs by being very sensitive and selective in identifying species to the lowest taxonomic level. Nonetheless this identification is dependent on what is deposit in the searched Database. Despite this fact, molecular biology proved to be an excellent approach for complementing the classical microcopy method. There are several molecular techniques used to detect phytoplankton species in water samples (McNamee et al. 2013). DNA hybridization assays (Goodwin et al. 2005), namely, Fluorescent Hybridization Assays (FISH) and Sandwich Hybridization Assays (SHA) (Guillou et al. 2002, Bertozzini et al. 2005, Goodwin et al. 2005, Antonella & Luca 2013) which can use whole cells that can be counted manually. In terms of Polymerase Chain Reaction there is for example, the real-time PCR or quantitative PCR (qPCR), which can provide estimation of microalgae cells concentration (Penna S.; Perini, F.; Bastianni, M.; Riccardi, E.; Pigozzi, S.; Scardi, M. 2012) and the conventional PCR that only gives the presence/ absence response. qPCR is capable of identifying and quantifying phytoplankton species, depending on the nucleic acid purification protocol, target gene, quantification of the standard curve and type of chemical detection method (Antonella & Luca 2013). Both conventional and qPCR, if they are meant to be used to substitute or complement classical microscopy techniques, they must be promising in both, identifying and quantifying phytoplankton in field samples. In that manner, nuclear ribosomal DNA, has been extensively used as a molecular target for the identification HAB species and other phytoplankton species. The most employed regions are the internal transcribed spacer (ITS/ ITS1-5,8S-ITS2) (Hubbard et al. 2008), the small subunit of ribosomal RNA (SSU/18S) (Metfies et al. 2005) and the large subunit of ribosomal RNA (LSU/28S) (McDonald D.; Zingone, A. 2007).

Anyhow molecular techniques are not always a straight forward protocol. It is known that Lugol’s solution and salts could act as a PCR inhibitor (Godhe et al. 2002, Galluzzi et al. 2004, Schrader et al. 2012), as well as humic acids, heavy metals and phenolic compounds that are also present in environmental samples (Park et al. 2009). In addition, small DNA amounts could also compromise DNA amplification. Samples from oceanic water can have for example only dozens of cells per milliliter of large dinoflagellates or even less. Monitoring phytoplankton samples, have usually these constraints. Biomass constrain can be overcome in bloom events or phytoplankton samples collected with a net. That said, an efficient genomic DNA extraction procedure is crucial for downstream steps. The efficiency of the genomic DNA extraction could determine the presence or absence of a phytoplankton species and even the quantification of that species in the sample. For that reason, it is of major importance to find out which method is more efficient in terms of quantity and quality for genomic DNA extraction. In addition, it is also determined the efficiency of the enzyme used in the PCR technique, once each DNA polymerase is developed aiming specific applications, such as sensitivity to amplify an exact fragment of interest and the ability to work under PCR inhibitors. Define the best kit to extract DNA from phytoplankton samples and the DNA polymerase able to amplify the target species of HABs, even if in low amounts, in natural samples is the objective of this work.

Previously, other works (Bertozzini et al. 2005, Simonelli et al. 2009, Eland et al. 2012) also compare comercial kits for phytoplankton DNA extraction, or even *Taqs* for animal samples (Arezi et al. 2003, Purzycka et al. 2006, Miura et al. 2013, Joana & Azevedo 2015, Tahir & Yaqoob 2016). This updated work goes in deep comparing more DNA extraction kits (12) and seven DNA polymerases for phytoplankton samples, to find out the best option for phytoplankton monitoring samples (PMS). In this way, to our knowledge this is the first work, comparing DNA polymerases activity for phytoplankton monitoring samples. In this work we choose as our species of interest *Pseudo-nitzschia sp*. This diatom occurs frequently in our coast and it is associated to Amnesic Shellfish Poisoning (ASP) syndrome. It often forms dense blooms that can be multi-specific with cryptic species. For that purpose, it was extracted genomic DNA from field samples known to have *Pseudo-nitzschia* species, fixed with Lugol’s solution and from *Pseudo-nitzschia sp*. cultures, fixed with Lugol’s solution and without fixation. The fixed cultures were used as positive controls, while non-fixed cultures were used to infer about the negative PCR interference by Lugol’s solution.

## Methods

### Sample preparation

Phytoplankton monitoring samples consist of sea water samples collected in the high tide, fixed with 0,4% (v/v) Lugol’s solution. Culture samples were obtained by the Phytoplankton Laboratory at IPMA through the isolation of *Pseudo-nitzschia* species collected from the Portuguese coast, followed by growth in f/2 enriched with silica medium (Guillard 1975), at 19°C, with 120μmolm^−2^s^−1^ on a 12:12 L:D cycle, for 5 days, in the Marine … (Marble) of IPMA. For total DNA extraction from cultures, it was used 50ml of sample, approximately 30×10^6^ cells. Culture samples were treated with and without the addition of 0,4% (v/v) Lugol’s solution. For total DNA extraction from monitoring samples it was used 200ml of sample, approximately 17×10^−4^ cells. All samples were centrifuged at 3500g for 10min (Dyhrman et al. 2006, Galluzzi et al. 2009). Thereafter, the supernatants were discarded, and the pellets stored at −20°C, until used. With the aim of maintaining the same conditions for all the experiments, we have chosen to freeze both fixed and non-fixed samples. In addition, this step could facilitate cell lysis.

### DNA extraction

The cell wall rupture, is the first step in plant DNA extraction protocols and if is not complete it compromises the subsequent steps of DNA extraction and amplification (Huang et al. 2000). Twelve different DNA extraction methods were tested. In Table 1, Arabic numbers designate the different DNA extraction methods used. Three commercial kits were tested only with chemical digestion (*3*, *4*, *6*), five with mechanical and chemical digestion (*1*, *5*, *7*-*9*), two fully automated (*8* and *9*), two that use the Direct PCR protocol (*10* and *11*) and finally the traditional phenol: chloroform method (*12*).

**Table 1.**
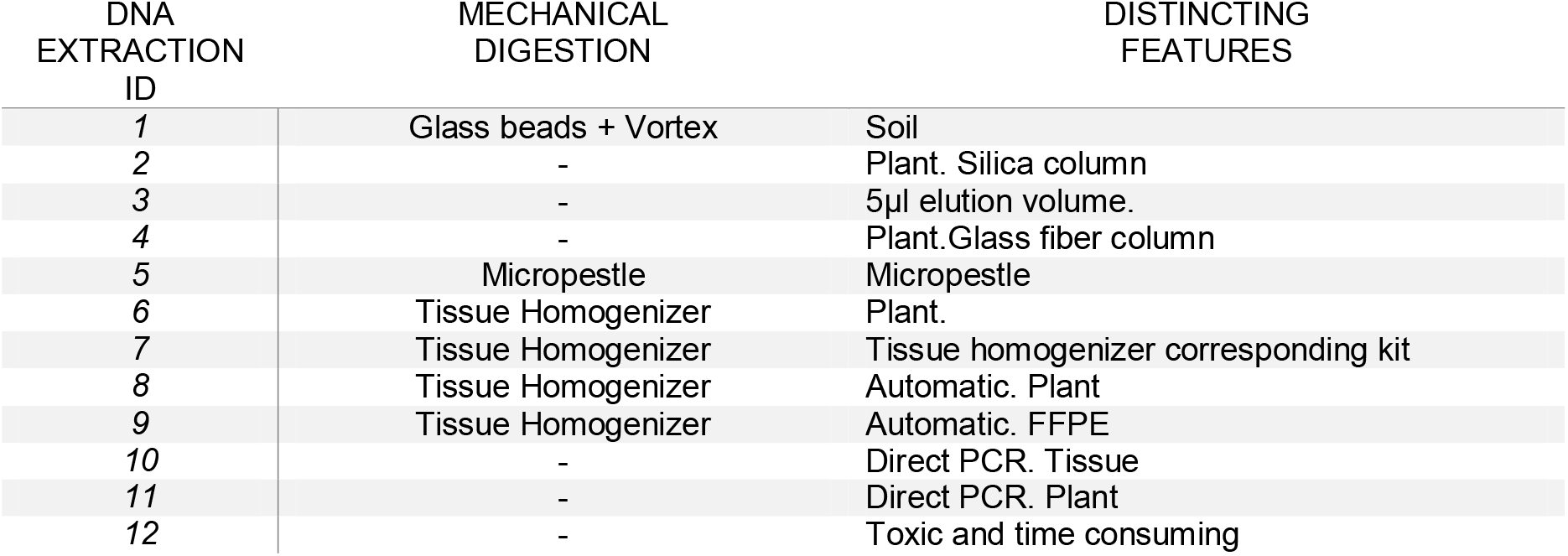
DNA extraction method used

All kits were used with the manufacture’s recommendation, with the following exception. Protocols from kits *2* and *4* advise to grind the sample in liquid nitrogen. However, the kits were manufactured for plant DNA extraction, which does not have the low biomass constrainer of microalgae. In this work, it was chosen to exclude this step once monitoring samples have low concentration of biological material. For example, for plant DNA extraction it can be used 50mg of biomass. If this procedure was performed, a biomass loss would exist from the adhesion of the sample to the mortar and pestle, resulting in a consequent DNA loss.

Summarizing, throughout this work, four types of methods/equipment of mechanical digestion, were used. Kit 1 had an agitation of a microtube with glass beads in a vortex (Vortex-Genie®). This process makes an agitation of 2700 rpm. Kit 5 used a manual micropestle, used inside the microtube. Kit 7 made a mechanical digestion of 6000 rpm for 40sec in a mechanical homogenizer. And, finally, Kits 6, 8 and 9, it was tested the Minilys® personal homogenizer (Bertin, corp.) with 5000 rpm for 15sec.

The Phenol: chloroform method was lead according to Band-Schmidt et al. (2003) with some modifications. Briefly CTAB lysis buffer (Panreac, AppliChem) with 0,2% β-mercaptoethanol (Sigma:Aldrich), was added to the sample and left to digest 1hour at 60°C. Then a mixture of phenol: chloroform: isoamyl alcohol (25:24:1) (Sigma:Aldrich) was added, mixed and centrifugued at 4°C and thereafter the supernatant was discarded. Sodium acetate (Panreac, AppliChem) and 2.5 volumes of absolute ethanol (Sigma:Aldrich) was added to the pellet and incubated for 1hour at −80°C. For DNA precipitation, a centrifugation step of 15min at maximum speed was performed. Afterwards DNA was washed with 70% ethanol. A final centrifugation was performed, and the DNA was ressuspended in 50μl of TE.

The choice of the various kits relies on its different characteristics, and the purpose was to infer which of them were more appropriate to PMS. kits *1* and *2* are based in the spin column technology. The spin column technology is based on the solid phase extraction method, where the DNA binds to the solid phase (column of silica) while the contaminants are washed throughout the column. This procedure is conducted by a series of steps of solutions addition and centrifugations that culminates with the DNA elution from the column. The soil kit (1), can remove trace contaminants, that in this case could be appropriate for Lugol’s fixed samples. The plant kit (2) is capable of eliminating polysaccharides, phenolic compounds, and enzyme inhibitors from plant tissue lysates, which in this case could be appropriated as microalgae have plant features, namely polysaccharides (Rether et al. 1993). In the kit *3* the elution can be as little as 5 μL resulting in concentrated DNA, that could be very advantageous for monitoring samples that have low quantity of biomass. Kits *4* and *5* are from the same brand and based on the same column methodology, but varies in the target, plant and tissue respectively. In these kits the matrix that traps DNA is not silica, but instead is glass fiber. Kit *5* is appropriated for a variety of tissue, including FFPE tissues. This could be helpful with samples fixed with Lugol’s solution, once as paraffin, Lugol’s solution also inhibits PCR techniques. Kit 6 is another column-based kit that also ensures the removal of most PCR inhibitors, but from a different brand from the remaining. The kit *7* can be used apart from the mechanic homogenizer of the same brand, yet they recommended to be used together. Therefore, we test the pair kit/ homogenizer according to the manufacturer’s instructions. Concerning to kits *8* and *9*, they belong to an automated system for DNA extraction. One is for plants and the other for FFPE tissues. This system has as major advantage the obvious fully automation. Apart from that reason, it is emphasized its easy reproducibility and the minimum cross contamination. After a tissue homogenization, the equipment uses a magnetic bead separation technology, where a magnetic bead with a silica surface attaches de DNA, the bead then responds to a magnetic field when all the unbound material is washed and finally the DNA is eluted from the bead.

The Direct PCR kits (*10* and *11*) do not perform a true DNA extraction. DNA is released from cells with the adding of a reagent and the PCR is performed in that complex mixture. DNA polymerases from this kit are claimed to be able to execute its work in the presence of PCR inhibitors. The tube that carries the crude sample (monitoring sample or culture) is the same where is stored the template DNA. The process prior the PCR reaction happens very fast and if the DNA released is for a single use, the PCR reaction occurs in the same tube that carries the crude sample. Fig 4 represents the Direct PCR workflow.

Finally, the traditional phenol: chloroform (12) method was tested. This method can extract high amounts of DNA, nonetheless is time consuming, being easily contaminated and the exposition to dangerous chemicals is an evidence to the handlers.

Genomic DNA concentrations (ng/μl) and contamination inference (Abs260/280 ratio) were accessed by Qubit Fluorometric Quantitation (ThermoFisher Scientific) and NanoQ spectrophotometer (Optizen). Genomic DNA integrity was assessed by gel electrophoresis on 0,85% agarose gel. The performances of the twelve methods tested were compared in terms of DNA amount recovered, quality and amplificability. The *Taq* performance was evaluated by positive species identification by sequencing.

### Polymerase Chain Reaction

To compare DNA extraction methods

To infer the best DNA extraction kit, it was amplified a fragment of the LSU from *Pseudo-nitzschia* sp., with the primers pair Pseudo 5’/Pseudo 3’(Penna et al. 2007). It was used fixed and unfixed *Pseudo-nitzschia sp*. cultures and PMS. Primers were synthesized by Invitrogen™ and the amplifications were performed on a T100™ thermal cycler (Bio-Rad). Reaction tubes contained the volume and reaction reagents concentrations specified by the *Taq* used, as well as the cycling conditions. PCR products were resolved in 2% (w%v) agarose gels and run in 1% TBE buffer. Gels were visualized by GreenSafe Premium (Nzytech) under UV light. GRS Ladder 100bp (GRISP) was the marker used. In Fig. 1 it is shown the several PCR amplifications profile for the 12 procedures.

**Fig 1.**
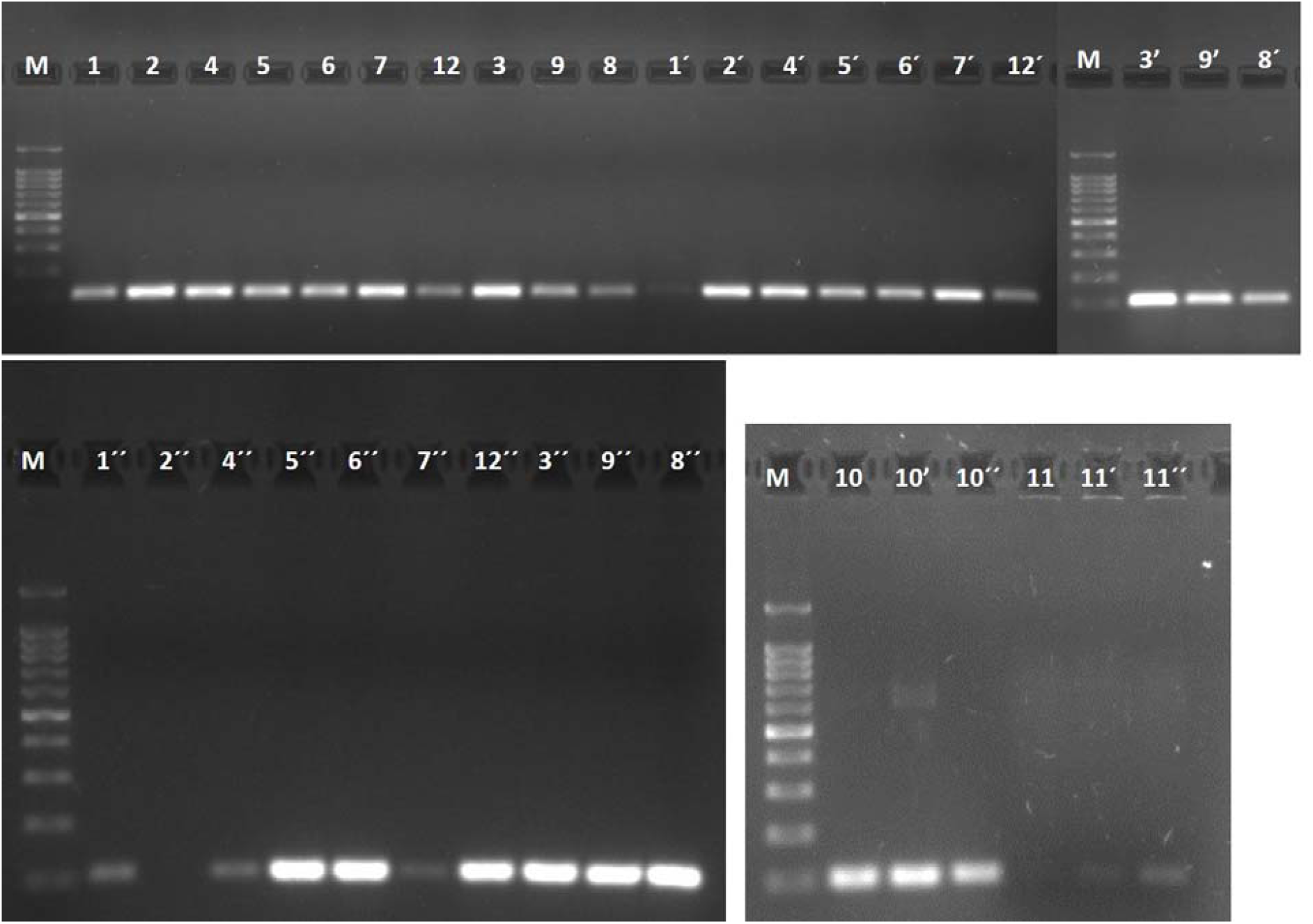
*Pseudo-nitzschia* LSU amplicon, from tested DNA extraction methods. Unfixed, fixed cultures and monitoring samples are represented by arabic numbers without a prime, with a prime or with a double prime, respectively. The correspondence between the numbers and the kit characteristics, are in table 1. *M* is the marker (GRS Ladder 100bp (GRISP).

### To compare *Taqs* performances

The various *Taqs* were tested against the same DNA sample, culture or PMS, both extracted with kit 2, and were used according to the manufacturer’s instructions. Table 2 designates by roman numbers the *Taq*s used.

**Table 2.** Distinct characteristics of the *Taqs* used

| *Taq* ID | Special Features |
| --- | --- |
| *I* | Direct PCR master mix for animal tissue |
| *II* | Direct PCR master mix for plant tissue |
| *III* | PCR master mix bead technology |
| *IV* | Hot start *Taq* w/ master mix |
| *V* | High sensitivity *Taq* w/ master mix |
| *VI* | Hot start *Taq* |
| *VII* | High fidelity *Taq* |

In this work, it was tested two Direct PCR mixes (*I* and *II*); three master mixes (*III, IV, V*) where one (*III*) is supplied in a bead format and the reagents are freeze-dried; and two stand-alone enzymes (*VI* and *VII*). *Taq I* and *II* correspond to Kit *10* and *11* respectively, as they cannot be dissociated from the DNA extraction process included in the Direct PCR protocol.

Among Direct PCR protocols, we have chosen to test one for tissue samples (*I*) and other for plant samples (*II*). Direct PCR *Taqs* are said to be very robust and able to perform PCR in the presence of PCR inhibitors. They are also very sensitive and able to amplify very low amount of template DNA. These features make these enzymes, very appropriate for PMS. Master mixes have the advantages of less pipetting errors, faster PCR mix preparation and have the option to exclude the loading dye when loading the samples in the electrophoretic gel, saving time and effort. In this category (master mixes), a PCR bead protocol was tested. PCR beads (*III*) assert to represent minor risks of pipetting errors and stability for storage at room temperature. Another master mix used (*IV*) declared to have more than 100x *Taq* fidelity for robust amplification of difficult targets and a hot-start technology. Hot start technology means least activity at room temperature, which avoids the extension of unspecific products from the primers or primer-dimers, and in that sense, produce a higher specificity of DNA amplification. By last, *Taq V* claim to have a higher sensitivity compared to conventional polymerases. In opposition to PCR master mixes, stand-alone *Taqs* although could be subjected to higher pipetting errors, have the advantage to allow MgCl2 concentration adjustments for PCR optimization. In this case it was tested two *Taqs* (*VI* and *VII*) with hot-start like capacity, which allows easiness and convenient *Taq* handling.

For *Taq* performance evaluation, the primers pair Pseudo 5’/Pseudo 3’(Penna et al. 2007) were used for *Pseudo-nitzschia sp*. fixed cultures and PMS. The sample used was the same for all the tested *Taqs*, DNA extracted from kit *2*, except for Direct PCR protocols that used the DNA extracted in the first step of the protocol. The reaction was performed on the Thermal Cycler referred previously. The reaction tubes contained the volume and reaction reagents concentrations specified by the *Taq* manufactures, as well as the cycling conditions. The electrophoresis was performed as previously described. In Fig. 2 it is shown the different PCR amplifications for the seven tested *Taqs*.

**Figure 2.**
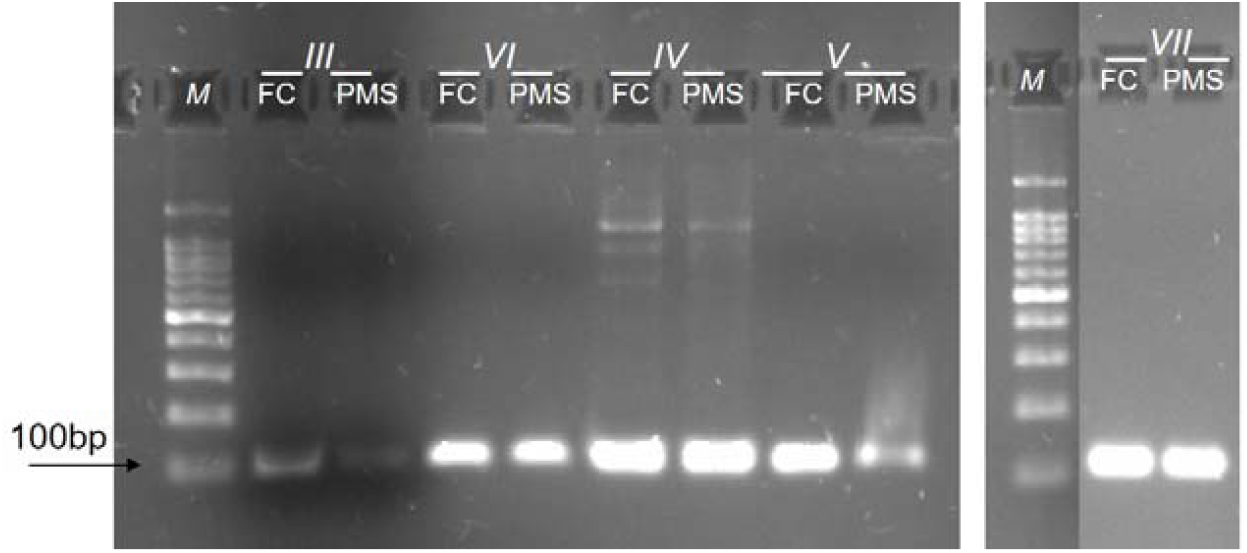
Taq performance assessment. *Pseudo-nitzschia* sp. fixed cultures (FC) and phytoplankton monitoring samples (PMS) were tested against 7 *Taqs*. The correspondence between *Taq* number and *Taq* characteristics is on Table 2. *M* is the marker (GRS Ladder 100bp (GRISP).

**Figure 4.**
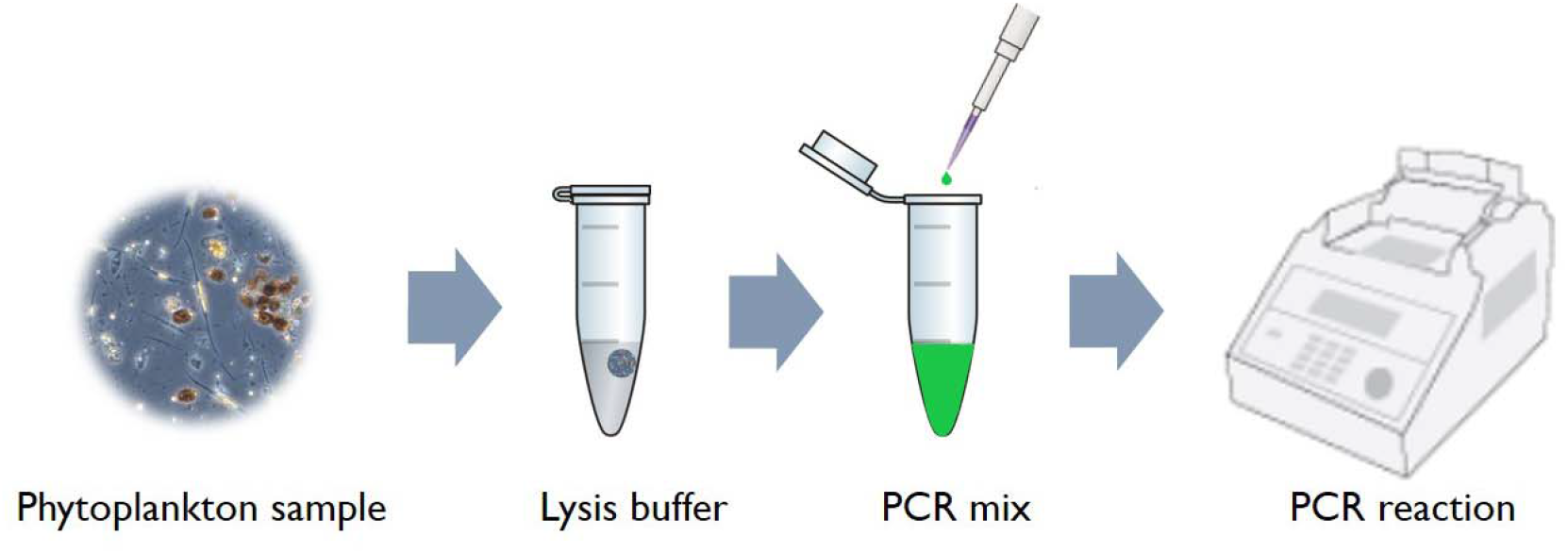
Direct PCR workflow.

## 2. Results

### PCR to compare DNA extraction methods

Results of quantity, purity and integrity of total genomic DNA extracted from unfixed, fixed cultures and monitoring samples were as expected (data not showed).

Fig. 1 shows the amplificability of the genomic DNA extracted, evaluated under a PCR with a species-specific primer pair for *Pseudo-nitzchia* sp. Fragments of the expected size (100bp) were obtained for all the samples. The fixative Lugol’s solution did not interfere with the PCR reaction. All tissues homogenizers used in kits *1*, *5*, *7*, *8* and *9*, not increase or decrease PCR yield compared to kits with only chemical digestion. Comparing Direct PCR protocols, Kit *10* retrieved higher yield compared to kit 11, regardless of kit 10 for fixed cultures amplified some unspecify product of higher molecular weight.

### PCR to compare *Taq* performance

All the tested *Taqs* could amplify *Pseudo-nitzschia* sp. from monitoring samples as well as *Pseudo-nitzschia* sp. cultures fixed with Lugol’s solution. See Fig. 1 for Taq *I* and *II* (kit *10* and *11* respectively) and Fig. 2 for the remaining (*III-VII*). Yet, *Taqs I*, *V*, *VI* and *VII* showed better results compared to the others (*II*, *III* and *IV*). *Taqs II* and *III*, showed slightly less yield of amplified product and *IV* showed amplification of unspecific products.

## 2. Discussion

In general, a laboratory which works with monitoring analysis, processes several samples at the same time. Therefore, time is crucial for having a replay to the final consumers. Direct PCR protocol with PCR master mix, was far from the fastest of all the methods tested and for that reason is one that should be considered.

In a general way, all methods worked well. The addition of Lugol’s solution didn’t have the negative interference as some authors report (Auinger et al. 2008). Otherwise, all the methods worked well for the samples with or without fixative. These results are also supported by others as Mäki and co-workers (2017). Some monitoring samples gave poor results possible because some inhibitory contaminants of the sample or the suitability of the kits. It can be discarded the variability of the initial DNA amount, once that was nearly the same for all the samples and the Lugol’s interference, based on the previous reaction where culture samples non-fixed *vs* fixed have the same output.

Several plant DNA extraction kits suggest the use of a mechanical tissue homogenizer, for complete fragmentation of the cell wall. Phytoplankton cells are similar to plant organisms in many ways, namely in the existence of a hard theca, for some dinoflagellates, or in the case of diatoms, as *Pseudo-nitzschia*, frustules. Thecae and frustules are rigid structures that protects cells and are made of cellulose or silica respectively. In that sense it would be expectable, that DNA extraction kits without this homogenization step, retrieved less DNA yield that would imply less PCR product. In fact, Yuan and coworkers (2015) used a bead-beating method that enhanced the final yield of DNA (highest as 2 folds) in comparison with the non-bead-beating method tested. However, no differences were encounter with different digestions. Even comparing with or without mechanical digestion or between mechanical digestion equipment. From this work, it was seen that the homogenization with glass beads in a vortex is simple, fast and adequate do phytoplankton cells. The homogenization with a micropestle, withdraw biomass, that is already few at the beginning of the procedure. Moreover, is time consuming and can lead to contamination. Finally, it was tested two types of mechanical homogenizer. One that the maximum speed is 5000 rpm (speed used) and the other that goes to 6800 rpm (6000 rpm used). It was verified that the minimum speed used is enough for fragment frustules and above this velocity, DNA can be shredded. As our amplicon is short (100bp) both velocity worked well, however if long fragments are to be amplified, the first option is the ideal.

Concerning to the kits performance particularly for monitoring samples, four kits had less PCR yield. Kit aiming soil samples (1) probably due to the type of sample required. Kit for plants with glass fiber column (4) and Direct PCR for plant cells (11). This fact can be due to the characteristics of the kit itself. Kit that used a high-speed tissue homogenizer (7) can possible be due to the excessive force applied for fragmenting the cells Previously to our work, others have also tested microalgae DNA extraction methods performance. One work tested six kits (Simonelli et al. 2009) and the other three kits (Eland et al. 2012). Both works tested soil, plant and tissue kits. In both, tissue kit gave the best result, in opposition to what it was expectable for plant kits. In our work, it was tested one soil kit, five plant kits and five tissue kits. Among those, two pair of kits (4&5 and 8&9) were from the same brand, varying in the target sample, plant and tissue respectively. In all the tested samples the results were good and similar, except for the kit 5 (tissue), that retrieved better results in the PMS. Comparing the direct PCR kits (10&11), of tissue and plant respectively, also the kit for tissue samples, gave higher yield of PCR. Nonetheless, in culture samples fixed with Lugol’s solution, appeared an unexpectable band of higher molecular weight, possible due to contamination in the Lugol’s solution. Hereupon, it can be concluded that from our work, the tissue kit worked well for phytoplankton PMS compared to plant or soil kits.

Concerning to the automated kits (8 and 9), previously, Bertozzini et al. (2005) compared four methods for extracting DNA from phytoplankton samples. Two methods based on super-paramagnetic beads and two without that technology. In our work, paramagnetic beads result as good as the other tested kits, in opposition to what was cited, that super-paramagnetic beads kit retrieved better results.

Kit 10, Direct PCR protocol with PCR master mix, gave the same good outputs as the conventional kits, with the advantage of processing the sample in significantly less time. Furthermore, processing the samples with less steps reduce possible contaminations, and the use of a master mix avoid pipetting errors and decrease time processing and thus retrieving more reproducible results. Beyond that, this kit was able to perform the PCR amplification with all the possible contaminants of the sample including the fixation agent. Lugol’s solution, with no need of a mechanic homogenizer. In addition, is a cost-effective solution, once although this kit is more expensive, it discards the use of a conventional DNA extraction Kit and it is only need this one product for extracting, PCR and gel loading, therefore equalizing the coasts. Therewithal the PCR run also takes less time, compared to conventional ones. In a laboratory that weekly processes more than 50 PMS, these features are major advantages, once these protocols can be operated by various technicians without losing reliability. This protocol also seems to have potential to be capable of amplifying phytoplankton species that appears in low number, sometimes in the dozens of cells per milliliter. Summing up, kit *10* showed to be adequate being sensitive, robust and fast, for processing phytoplankton monitoring samples.

Regarding to the *Taq* performance, all the tested *Taqs* could amplified PMS as well as fixed cultures, yet *Taq IV* amplified also nonspecific products for both samples. Excluding the factors of sample contamination and primers specificity, once they were equal to all samples, this result could be due the suitability of the template program to phytoplankton PMS, being necessary a temperature optimization, or more unlikely that we are standing before a *Taq*, that introduces more error in synthetizing the product.

## Conclusion

Marine phytoplankton monitoring samples are routinely analyzed for early warning to bloom toxicity and to support HPLC results for toxin accumulation in bivalves, that are endanger to humans. In that sense, microscopy and molecular biology go hand in hand, in phytoplankton identification, being the second gaining relevance. From our study, the differences observed between methods are not determinant to exclude a particular kit. Roughly, Lugol’s solution did not interfere negatively with PCR. There are kits available in the market that do not need an initial step of mechanical homogenization for an adequate DNA extraction, as is sometimes advised. However, if it is attended to use for magnifying DNA extraction, a mechanical homogenizer with beads with a medium force applied, such as 5000 rpm, is adequate for phytoplankton cells. Once little force does not work and by the other hand, to much force applied can shredded DNA. Concerning to *Taq* performance, as previously, none should be excluded, even if some are able to amplify with high or less yield. Apart from what was said, there is one kit that stands out. Direct PCR master mix for tissue samples, proved to have the best performance. It was easy to handle, very fast, with reproducibly results and sensitivity enough to perform a PCR reaction in a mixture with PCR contaminants and low DNA content.

